# LongAxis: a MATLAB-based program for 3D quantitative analysis of epithelial cell shape and orientation

**DOI:** 10.1101/684555

**Authors:** Keith R. Carney, Chase D. Bryan, Hannah B. Gordon, Kristen M. Kwan

## Abstract

Epithelial morphogenesis, a fundamental aspect of development, generates 3-dimensional tissue structures crucial for organ function. Underlying morphogenetic mechanisms are, in many cases, poorly understood, but mutations that perturb organ development can affect epithelial cell shape and orientation – difficult features to quantify in three dimensions. The basic structure of the eye is established via epithelial morphogenesis: in the embryonic optic cup, the retinal progenitor epithelium enwraps the lens. We previously found that loss of the extracellular matrix protein *laminin-alpha1* (*lama1*) led to mislocalization of apical polarity markers and apparent misorientation of retinal progenitors. We sought to visualize and quantify this phenotype, and determine whether loss of the apical polarity determinant *pard3* might rescue the phenotype. To this end, we developed LongAxis, a MATLAB-based program optimized for the retinal progenitor neuroepithelium. LongAxis facilitates 3-dimensional cell segmentation, visualization, and quantification of cell orientation and morphology. Using LongAxis, we find that retinal progenitors in the *lama1^−/−^* optic cup are misoriented and slightly less elongated. In the *lama1;MZpard3* double mutant, cells are still misoriented, but larger. Therefore, loss of *pard3* does not rescue loss of *lama1*, and in fact uncovers a novel cell size phenotype. LongAxis enables population-level visualization and quantification of retinal progenitor cell orientation and morphology. These results underscore the importance of visualizing and quantifying cell orientation and shape in three dimensions within the retina.

## Introduction

Organogenesis requires assembly of cells into precise 3-dimensional structures which are crucial for function. Disruptions to this morphogenetic process can lead to organ dysfunction, and are a common cause of birth defects. Cellular and molecular mechanisms governing organ morphogenesis are generally not well understood: many signals and pathways have been identified, often on the basis of genetic studies and mutant phenotypes. Despite this, analyzing genetic interactions and dissecting how different factors impact morphogenesis has been a challenge, since, in many cases, it has not been trivial to visualize and quantify phenotypes in three dimensions.

The vertebrate eye forms via a complex morphogenetic process, during which the optic vesicle, an outpocketing of the forebrain, undergoes cell and tissue movements to become the optic cup, in which the hemispherical retina enwraps the lens. At the end of optic cup morphogenesis, the retinal epithelium is comprised of progenitor cells which are elongated and oriented toward the lens. In fish, mouse, and chick, genetic screens, candidate approaches, and conditional genetic studies have identified factors involved in optic cup tissue organization (Adler and Canto-Soler, 2007; Bazin-Lopez et al., 2015; Chow and Lang, 2001; Fuhrmann, 2010; Martinez-Morales and Wittbrodt, 2009; Yang, 2004). One key factor governing optic cup tissue organization and morphogenesis is the extracellular matrix, a complex proteinaceous layer that surrounds epithelial tissues and provides polarity, survival, and signaling cues (Adams and Watt, 1993; Daley and Yamada, 2013; Frisch and Francis, 1994; Juliano et al., 2004; Martin-Belmonte and Mostov, 2008). It has long been known that a complex extracellular matrix layer surrounds the nascent developing eye of all vertebrate species examined to date (Hendrix and Zwaan, 1975; Hilfer and Randolph, 1993; Kwan, 2014; Parmigiani and McAvoy, 1984; Peterson et al., 1995; Svoboda and O’Shea, 1987; Tuckett and Morriss-Kay, 1986; Wakely, 1977; Webster et al., 1984), and functional roles for specific extracellular matrix molecules in early eye development are starting to be resolved using molecular genetic approaches (Bryan et al., 2016; Hayes et al., 2012; Huang et al., 2011; Lee and Gross, 2007; Semina et al., 2006). We previously found that loss of *laminin-alpha1* (*lama1*) results in disruption of tissue polarity and cellular disorganization within the retinal epithelium of the zebrafish optic cup (Bryan et al., 2016). At the single-cell level, retinal progenitors appeared misoriented, although this seemed variable between individual mutant embryos and was largely inferred by scanning through volume data (z-stacks acquired by confocal microscopy). In addition to the tissue disorganization defect, the *lama1* mutant displayed ectopic localization of the apical marker pard3 at inappropriate locations, including what would normally be the basal surface of the optic cup. We wondered whether the establishment of ectopic apical surfaces might cause the disorganization phenotype, and whether removal of the apical determinant pard3 could rescue it.

Although we had questions, we lacked the methodology to adequately and quantitatively analyze such phenotypes. We were not previously able to visualize or quantify cell orientation in 3-dimensions, phenotypic variability between embryos, nor how changes in cell shape or volume might contribute to mutant phenotypes. With these goals in mind, we have developed LongAxis, a MATLAB-based program which allows us to qualitatively and quantitatively assay multiple aspects of cell morphology and organization, optimized for the developing retina. Using a combination of automated segmentation and refinement (or filtering) via user selections to remove outliers and incompletely segmented cells, we can visualize and analyze cell orientation and shape in 3-dimensions throughout the tissue. Cell orientation, length, length/width ratio, and cell volume can be calculated for thousands of cells simultaneously; these features can be displayed in the intuitively simple “urchin plot”, which conveys the cell’s extent of elongation (length/width ratio) and orientation.

Using LongAxis, we finally resolved questions regarding the *lama1* mutant optic cup phenotype, including how cell orientation and morphology are quantitatively affected, and whether genetic removal of the apical polarity determinant *pard3* is able to rescue it. We find that in the *lama1* mutant optic cup, retinal progenitors are indeed misoriented, and that misoriented cells cluster together in domains. Cells are shorter and less elongated, but not smaller than wild type cells. In the *lama1;MZpard3* double mutant, retinal progenitors are still misoriented, and we uncover a cellular-level phenotype: cells are larger than either wild type or *lama1* single mutants. Therefore, loss of *pard3* does not rescue the *lama1* mutant tissue organization phenotype. Importantly, rather than 2-dimensional measurements in a small number of sparsely labeled cells, LongAxis allows us to discover population-level alterations in cell morphology and organization, and underscores the importance of quantitative analysis of cellular level phenotypes.

## Results

### Pipeline for 3-dimensional cell segmentation

Our goal is to understand the molecular basis of cell and tissue organization within the embryonic optic cup. Although many factors have been identified as playing a role in this process, our analysis has largely been limited to 2-dimensional analysis of a small sampling of cells. Dissecting genetic interactions and mechanisms would ideally be carried out by quantitatively evaluating cell orientation and morphology throughout the retinal progenitor cell population. To this end, we developed LongAxis, a program to facilitate visualization and quantification of cell morphology within the zebrafish optic cup.

The goal of this software is accurate single cell segmentation and automated quantitative analysis of cell shape and orientation within the context of the tissue, therefore, a crucial initial optimization step is obtaining image data of adequate quality. To avoid distortion and changes in volume that accompany tissue fixation, we imaged live embryos at optic cup stage (24 hours post fertilization (hpf) in zebrafish), in which all membranes were labeled using EGFP-CAAX. Images were acquired at 1024×1024 resolution, and we aimed for adequate axial sampling without photobleaching, deciding upon a voxel size of 0.21×0.21×0.42 μm (x:y:z ratio of 1:1:2; Movie 1).

In LongAxis, cell segmentation begins with eight steps of 2D processing applied to every slice, with the goal of enhancing boundaries (Figure 1B; see also Methods, LongAxis MATLAB code). The processing steps outlined here are optimized for our specific data sets, the goal being to visualize and analyze retinal epithelial cells. In our experience, the key step was to correct for variations in signal (i.e. some regions of membrane around any particular cell might be brighter or dimmer than others) in order to ensure that the cells were segmented along membrane boundaries accurately. Once 2D processing has been carried out, the user selects the 3D volume of interest within the image data for 3D segmentation and rendering. In our case, this focuses our analysis on the retinal epithelium and excludes cells outside, such as prospective brain, lens, and overlying ectoderm. Once the subvolume of interest has been selected, 3D processing functions are applied to enhance and connect boundaries across slices (Figure 1C). This initially yields 3D cell segmentation throughout the volume data (Figure 1D, E). The 3D cell segmentation is then applied within the user-selected subvolume, leaving only cells within the region of interest (Figure 1F-H). Segmentation can be examined in small volume regions for visual validation at this stage (Figure 1I-J; Movies 2, 3).

**Figure 1.**
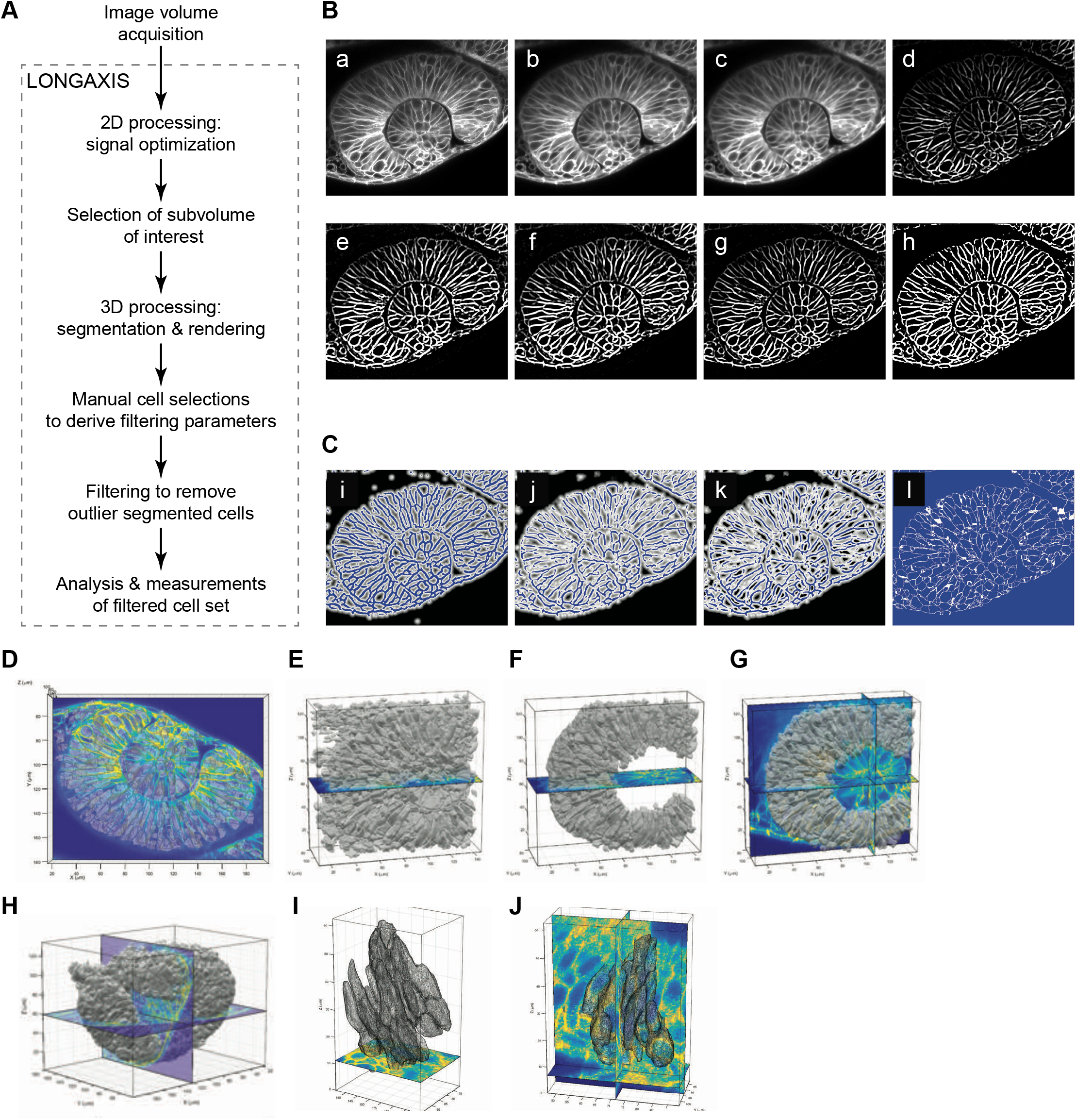
LongAxis workflow, image processing, and segmentation.

Following this, the set of segmented cells is refined: cell segmentation needs to be validated, and unwanted cells, particularly those in which segmentation failed, are removed from the data set. To this end, a process of “informed filtering” is carried out (Figure 1A). The basic idea is to validate cell shapes in a manner unbiased with respect to the orientation of the cell; assaying changes in cell orientation is a major goal of this software. To carry out filtering, an expert user (i.e. someone experienced with looking at these data) views 3-dimensional segmented cell shapes (away from the image data), and manually validates cells which appear to have a retinal epithelial morphology. If need be, the user can cross-check the position of the cell to ensure that cells within the retinal epithelium are being selected, or the user can also check the cell rendering against the original image data. The ability to cross-check may be useful in cases (e.g. mutants) where cell morphologies could be dramatically altered, but again, the basic idea is to carry out these selections in an unbiased manner with respect to position and orientation of the cell within the tissue.

Once the user has selected cells, the user-selected data set is analyzed: minimum and maximum values for cell volume, length, and length/width ratio are derived. These minimum/maximum (min/max) values for these three criteria are applied to refine and filter the entire data set (all segmented cells in the region of interest); this process thereby excludes “outlier” cells with respect to these three specific criteria. The filtered cell set, which represents all retinal epithelial cells (selected in an unbiased manner) is used for quantitative analysis of cell morphology and orientation and 3-dimensional visualization.

### LongAxis analysis and outputs

Once cells are segmented, a variety of outputs can be acquired, including 3-dimensional visualization of cell shape, and quantitative outputs including cell length, cell width, length/width ratio (a metric of how elongated the cell is), and cell orientation (Figure 2A-D; Movie 4). Cell orientation is quantified within the 3-dimensional tissue by deriving the cell convergence point: the average of all midpoints of closest approach for all cell orientation vector pairs (Figure 2E; Movie 5). This was empirically derived for each embryo independently: we found that existing landmarks (e.g. the lens center of mass) incurred too much variability between embryos, as lens shape, size, and even position can vary slightly with respect to the retinal epithelium. Once the vector convergence point is obtained, the long axis, which is derived from the ellipsoid fit, is used to calculate an angle of deviation (or deflection) from that convergence point for each cell (Figure 2F, marked by red asterisk). In addition to the quantitative output, angles of deviation and length/width ratio can all be represented in an “urchin plot”, a 3-dimensional visual representation of cell orientation and shape within the tissue (Figure 2G). In the urchin plot, the angle of deviation is represented by a heat map, in which close adherence to the expected angle is coded in bluer colors, and significant deviation is encoded by warmer colors. The length of the vector is proportional to the length/width ratio of the cell, to represent one aspect of cell shape.

**Figure 2.**
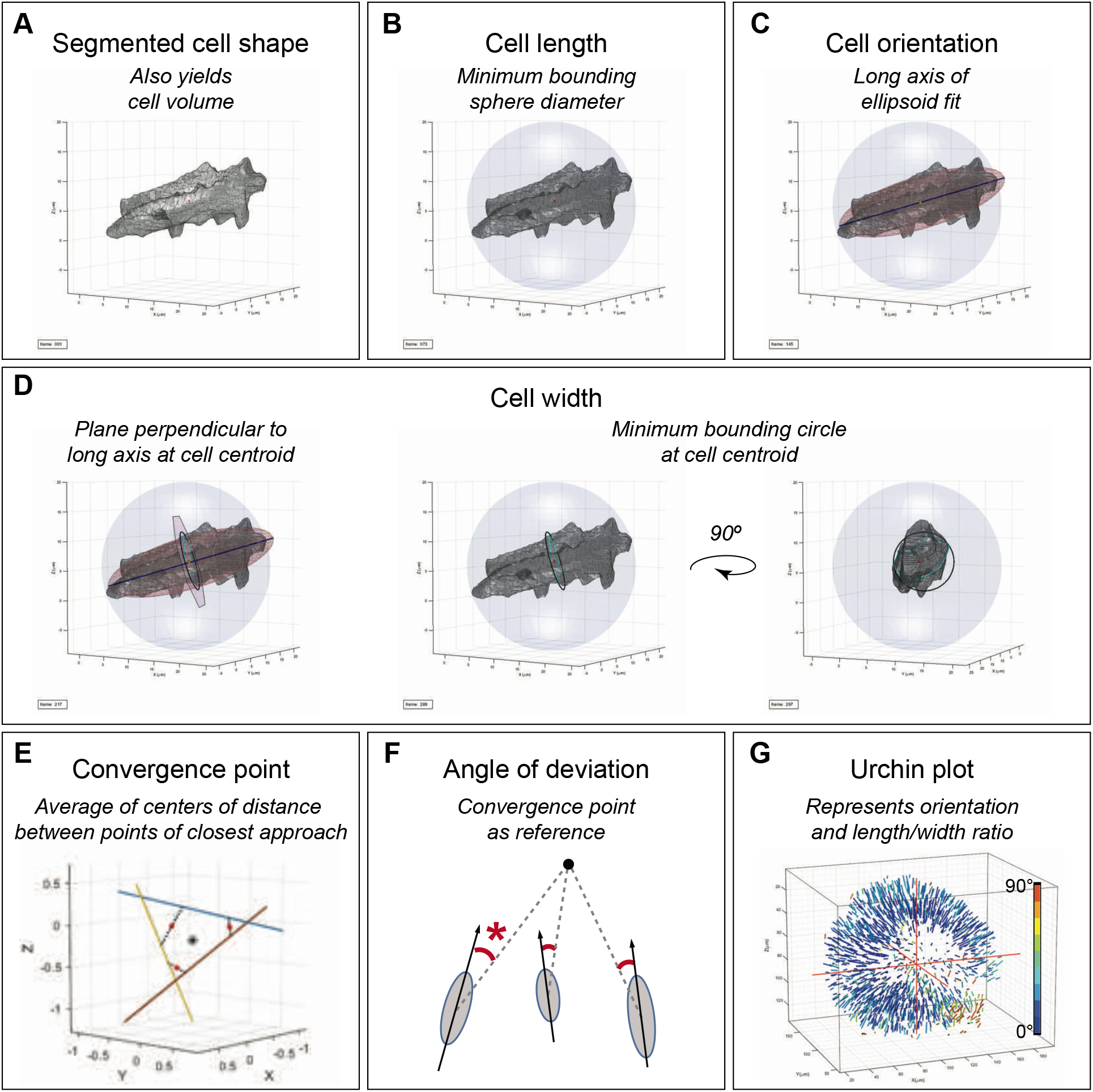
LongAxis data analysis and outputs.

### Validating segmentation and filtering

To determine how well the workflow performs, segmentation and filtering validation steps were carried out on three independent subregions of the image volume data (one example in Figure 3A). First, because segmented cell shapes were initially viewed in an isolated manner, away from the image data, we visually examined all segmented cells in each subregion against the original image data. We used xy, xz, and yz cutaways to evaluate how well the segmentation matched the membrane signal, including whether the process correctly segmented single cells. Segmentation accuracy for all cells was scored manually (by a user) on a scale of 1-5, with 1-4 corresponding to how well the segmentation matched the membrane boundaries in the image data (1 = 90-100% matching boundaries; 2 = 70-90%; 3 = 50-70%; 4 = <50%), and a score of 5 representing unsuccessful segmentation resulting in fused cells. Despite presence of some variability in rendering quality, cell orientation was largely unaffected for cells in categories 1-3. The proportions of cells in each category is shown in Figure 3B.

**Figure 3.**
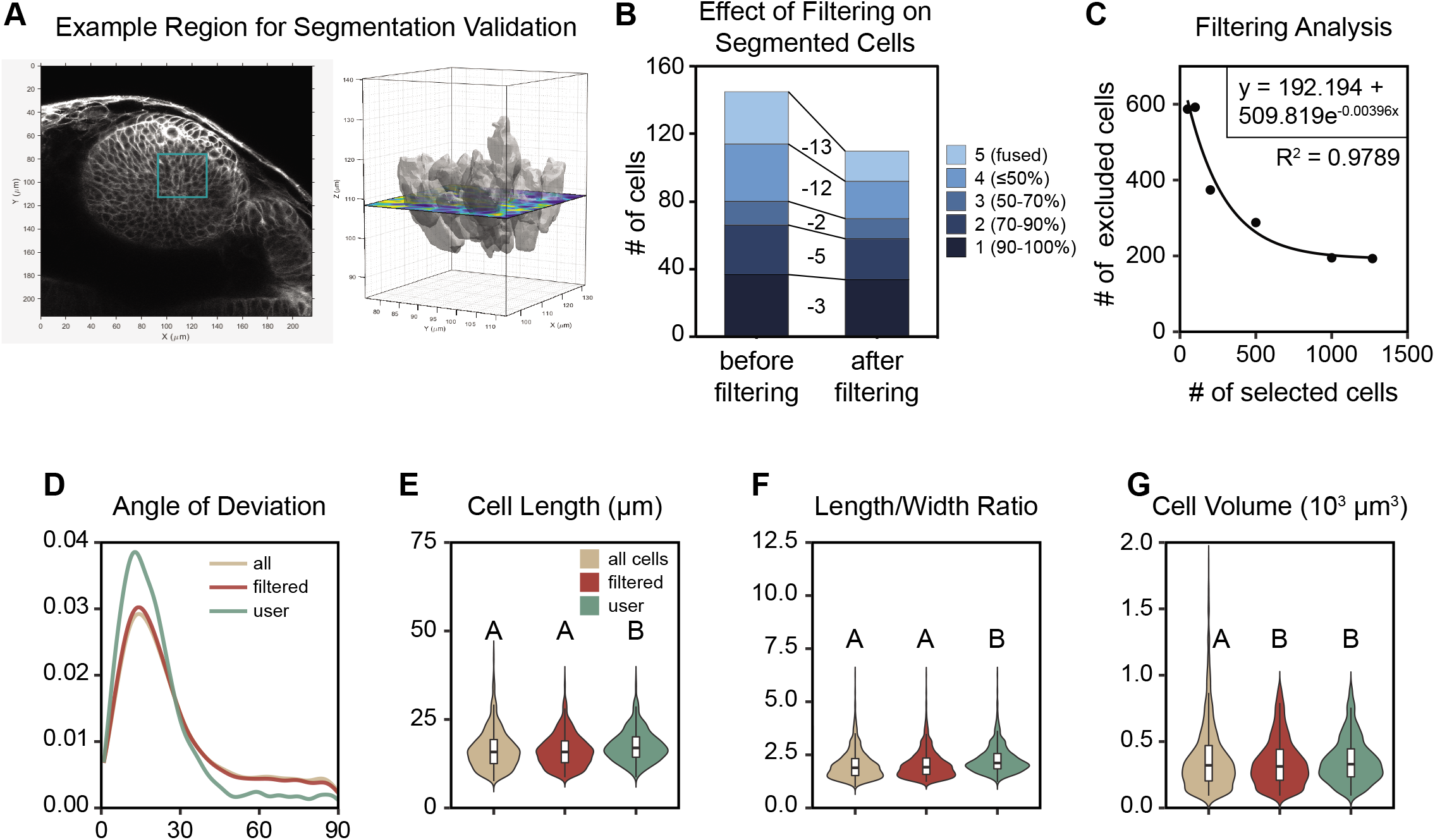
Filtering Analysis and Validation

We then asked how filtering (using parameters derived from user selections for cell volume, cell length, and length/width ratio) affected the number of cells in each group. We examined the subset of cells that passed the filtering criteria, and we found that indeed, although filtering is not perfect, poorly segmented (class 4) and fused cells (class 5) are preferentially removed from the filtered data set (Figure 3B; number of cells removed from each class 1-5, in order: 3, 5, 2, 12, 13). These analyses suggest that the segmentation identifies cells in 3-dimensions and filtering helps to remove unsuccessfully segmented cells, leaving us with a data set appropriate for population-level quantitative analysis.

### Determining filter parameters and the size of the user-selected data set

Accurate filtering relies on having a set of cells selected by an expert user; filtering parameters are derived from this user-selected cell set. How many cells does the user need to choose to generate reliable filtering parameters? We tested this in 4 different wild type embryos by examining the relationship between number of cells selected and number of cells filtered out, the rationale being that as the number of selected cells increases, more reliable filter parameters will be generated. This, however, only works up to a point at which selecting more cells has no more benefit; the user set will have already captured the full range of appropriately segmented cells. We find that the relationship between number of cells selected and number of cells filtered out obeys exponential decay (Figure 3C; Figure S1A); deriving the equation to describe this graph allows us to easily calculate the number of cells which need to be selected to carry out filtering (using the mean lifetime equation τ = λ^−1^, where λ is the decay rate and τ represents the mean lifetime, or here, the average number of selections it takes to remove a cell). For the 4 wild type embryos examined, although substantial numbers of cells were manually selected by the user (1269, 1788, 2420, and 1582, respectively), significantly fewer cells (using the mean lifetime equation to solve for τ: 253, 119, 120, and 61, respectively) needed to be selected in order to exclude the inappropriately segmented cells without inappropriately removing correctly segmented cells (Figure 3C; Figure S1A). While the number of user-selected cells necessary for adequate filtering needs only to be a small proportion of the total number of cells, filtering quality clearly increases with more user-selected and validated cells. In addition, the derived equation reveals that there is a minimum of cells that will be excluded in each wild type embryo (using the exponential decay equation (see Methods) and solving for *y_f_*: 192, 132, 200, and 515, respectively); based on our manual validation, these are likely to represent poorly segmented and fused cells. We think the variability in this number between embryos is due to variation in image quality, which will affect the success of the segmentation process.

This post hoc analysis reveals that there is not one single baseline number of cells for a user to select, however, the software is simple to use, and selecting a few hundred cells will likely yield high quality filtering information necessary to remove unwanted cells.

### Filtering poorly segmented cells does not change the data set

Given that filtering does change the number of cells being used for quantitative analysis (Table 1), we asked how it might alter, at the population level, the quantitative measurements of interest: angle of deviation, cell length, length/width ratio, and volume (Figure 3D-G; Figure S1B-E). We find that in the cases of angle of deviation, cell length, and length/width ratio, the distributions of filtered cells are not altered from the original (full) set of segmented cells (Figure 3D-F; Figure S1B-D, letters in graphs (A, B) represent different statistical groups). In contrast, cell volume is changed such that the filtered set is not statistically different from the user selections (Figure 3G; Figure S1E), consistent with the idea that inappropriately segmented, and especially fused cells (which are larger) are removed from the filtered data set. Importantly, distributions of orientation angles do not change (Figure 3D; in 3 out of 4 wild type embryos, two-sample Kolmogorov-Smirnov tests show no significant difference between the set of all segmented cells and the filtered set), so filtering would not influence large-scale analysis of cell orientation and tissue organization. We conclude from these analyses that filtering works to preferentially remove outlier cells, without changing the population distribution of the data with respect to cell length, length/width ratio, and angle of deviation.

### Wild type embryos exhibit slight morphological variability between cell populations

With our new tool in hand, we first set out to examine multiple wild type embryos to determine the amount of variability we might detect between samples of the same genotype. Development in zebrafish is not deterministic, and we expect there to be some variability between individuals with respect to eye size and cell number. To examine this, four different wild type embryos were imaged and analyzed using the LongAxis pipeline, with filtering parameters and cell convergence points derived independently for each individual embryo (Figure 4A-A”’, Movie 6). In Movie 6, isosurfaces show highest density regions of midpoints of closest approach for all pairwise vector combinations, and black dot shows the derived convergence point (the average of all calculated midpoints) which was used for angle of deviation measurements. Urchin plots were generated to visualize cell orientation in a qualitative manner (Figure 4B-C”’, Movie 7), and at this level of resolution, the optic cups exhibit some variation in size and shape (including lens shape). Despite this, the cells (represented as vectors in the urchin plot) largely appear to be aligned toward the calculated convergence point (labeled as colors in the blue range in the heat map), with the reproducible exception of the optic fissure opening at the ventronasal side of the eye (Figure 4B-C”’, asterisks). These data are represented quantitatively as a density plot of angles of deviation (Figure 4D); the distributions of angles of deviation appear similar between the four embryos, with a peak ~20°. Quantitative analysis and comparison of these 4 wild type embryos reveals other notable features. First, the numbers of retinal epithelial cells vary between the optic cups (1824, 2413, 3752, and 2019, respectively; Table 1), but these numbers are in the same range as previously calculated using a completely independent method which relied on counting nuclei (Kwan et al., 2012). This serves as a convenient independent validation of our approaches. Next, the distributions of cell length and volume appear different, but fall only into two statistical groups (Figure 3E, G; mean length (μm): 16.26, 15.9, 16.29, 16.59; mean volume (μm^3^): 346.28, 305.28, 337.3, 341.05). There is no significant difference in cell length/width ratio between the four embryos (Figure 3F; mean length/width ratio: 2.06, 2.08, 2.09, 2.1). Taken together, these data indicate that the quantitative analysis can distinguish between individual embryos of the same genotype, due to normal phenotypic variability; therefore, multiple embryos must be used to compare different experimental conditions and genotypes.

**Figure 4.**
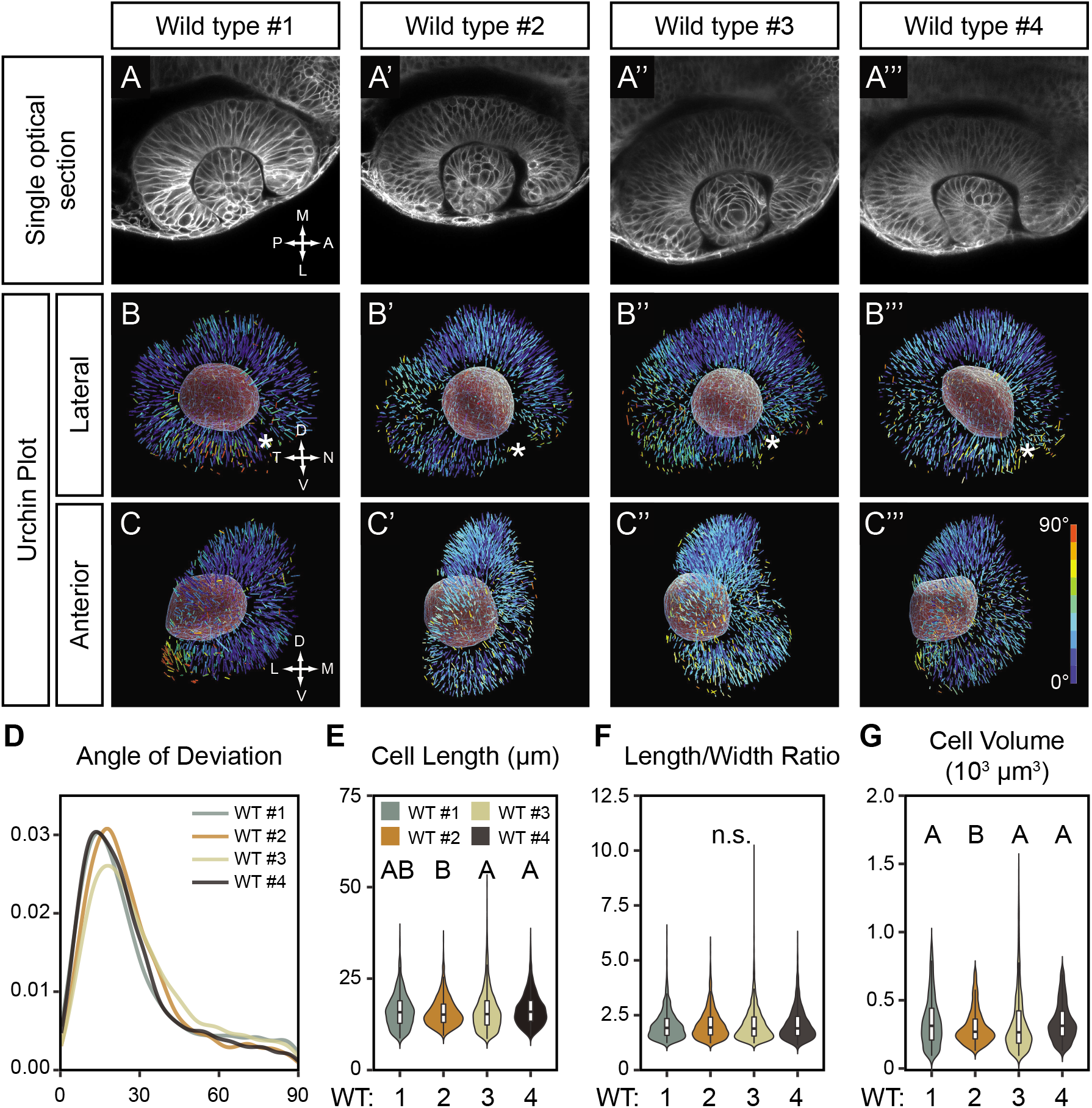

### Putting the software to the test: genetics of apicobasal polarity and tissue organization in the optic cup

Having determined that we could segment cells and carry out quantitative analysis on cell morphology and orientation, we turned our attention to the original biological question at hand. We previously demonstrated that loss of *lama1* leads to disruptions to epithelial polarity and apparent disorganization of the retinal progenitor epithelium (Bryan et al., 2016). Although we hypothesized that the cause of this phenotype was cell misorientation as opposed to gross changes in cell size or shape, we had no way at the time to visualize or quantitatively test this. In addition to the retinal disorganization, we found that tissue polarity is disrupted in *lama1* mutants: apical markers such as pard3 are mislocalized and even ectopically localized to subcellular locations that would, in a wild type embryo, be the basal surface. We wondered whether ectopic localization of apical determinants was the cause of the structural disorganization in the *lama1* mutant optic cup, and therefore, whether the *lama1* mutant phenotype might be rescued by genetic removal of *pard3*.

With LongAxis in hand, we set out to answer these questions. We generated double mutants for *lama1* and *pard3*, in which *pard3* was both maternally and zygotically lost (*lama1;MZpard3*), as pard3 is maternally loaded (Blasky et al., 2014). We compared wild type optic cups to the *lama1* single mutants and the *lama1;MZpard3* double mutants. When initially viewing single optical sections of all three genotypes (Figure 5A-A’’), the *lama1* single mutant exhibits the expected disorganized retinal epithelium with cells that appear cuboidal in cross section (Figure 5A’, Movie 8). The *lama1;MZpard3* optic cup initially appeared as though the disorganized retinal progenitor cell phenotype might be partially rescued (Figure 5A’’, Movie 9); in this optical section, some cells are elongated and oriented toward the lens.

**Figure 5.**
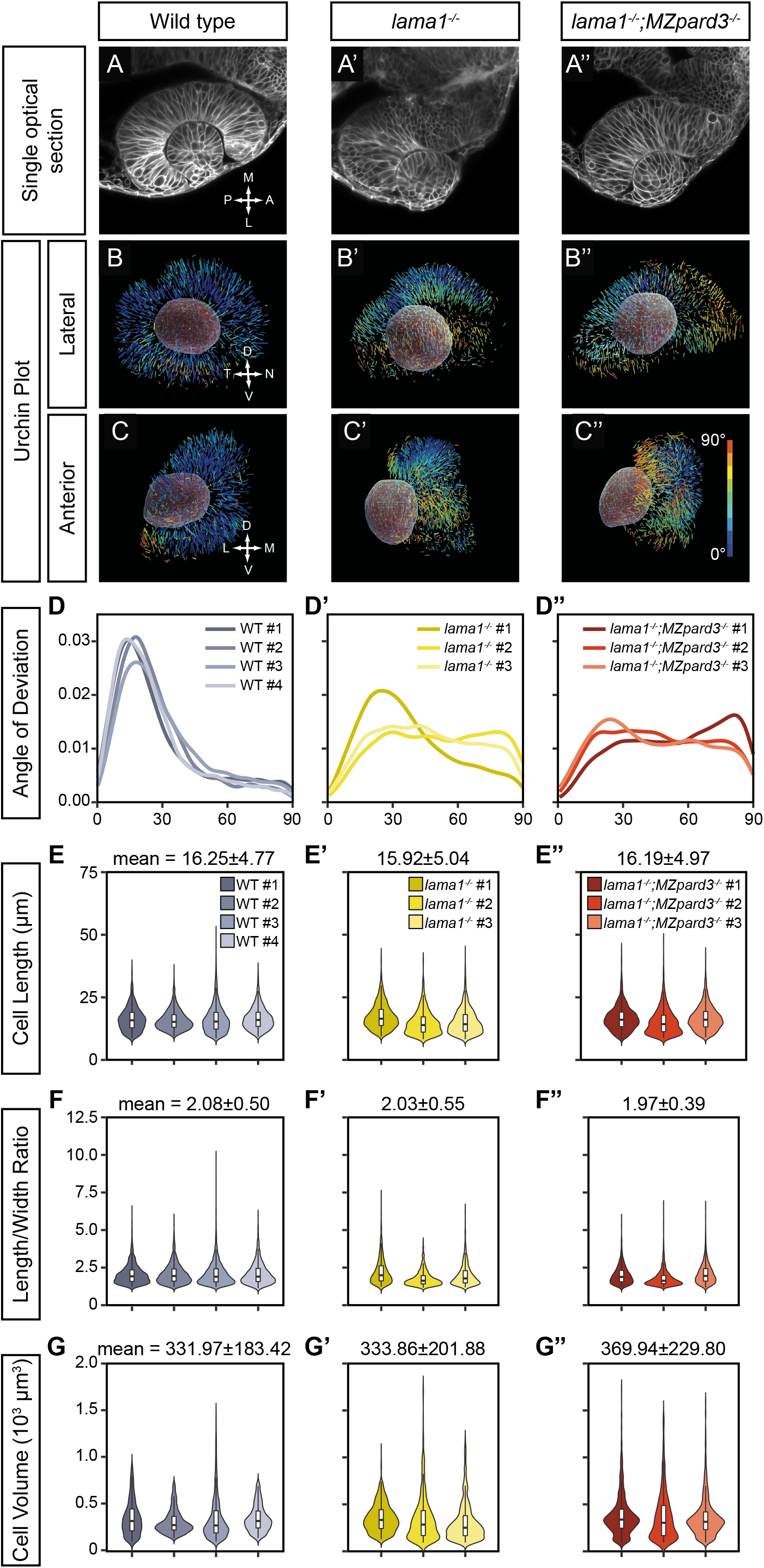

This, however, underscores the importance of our approach, as 3-dimensional visualization and quantification are necessary to actually resolve whether the phenotype is rescued or not. Three independent embryos of each genotype (*lama1^−/−^* and *lama1^−/−^;MZpard3^−/−^*) were imaged, processed through our LongAxis pipeline, and compared to the wild type optic cups. First, cell convergence points were derived (Movie 10) and urchin plots were generated to qualitatively visualize and compare cell orientation (Figure 5B-C’’; Movies 11, 12). Because of the heat map coding of the vectors, it is intuitively clear that significant regions of the optic cup in both *lama1* and *lama1;MZpard3* double mutant eyes are comprised of misoriented cells. Substantial patches of the urchin plots are populated by vectors in the red-orange-yellow range, indicating an angle of deviation >60°. Interestingly, misoriented cells were found clustered, as opposed to individually randomly scattered throughout the eye.

This is also represented quantitatively in the angle of deviation density plots (Figure 5D-D’’): the four wild type embryos all have a peak ~20°, indicating a small deviation from the convergence point, and a small trailing tail out beyond 60° (Figure 5D; Movie 7). In contrast, the *lama1* single mutants show a very different distribution in angles of deviation: in one embryo, there is a visible peak ~20°, similar to wild type embryos, but in the other two embryos, there is no clear peak, rather, angles of deviation are distributed more evenly from 20-90° (Figure 5D’; Movie 11). Similarly, all three *lama1;MZpard3* double mutants show an even distribution of angles of deviation from 20-90°, without a clear peak (Figure 5D”; Movie 12). This indicates that at the population level, cells are significantly misoriented in both the *lama1* single mutant and *lama1;MZpard3* double mutant optic cups.

In our previous work, we quantified morphology of a small number of cells to determine whether tissue disorganization might actually be caused by changes in cell length or length/width ratio. Assaying limited numbers of cells primarily in 2 dimensions, we found that retinal progenitor cell length was diminished, but length/width ratio was unaffected (Bryan et al., 2016). Although we had obtained a preliminary answer to our question, we wanted to determine if these trends held true with more thorough quantification of cell morphology across the population of retinal progenitors.

Using our LongAxis pipeline, we compared retinal progenitor cell length, length/width ratio, and volume at the population level, with >1000 cells per eye. First, in terms of cell length (Figure 5E-E’’), we find that *lama1* mutant retinal progenitor cells are shorter than wild type; this is consistent with our previous data (Bryan et al., 2016). In contrast, however, *lama1;MZpard3* double mutant retinal progenitor cell length is indistinguishable from wild type (wild type 16.25±4.77 μm; *lama1^−/−^* 15.92±5.04 μm; *lama1^−/−^;MZpard3^−/−^* 16.19±4.97 μm). Next, we examined length/width ratio: although our previous 2-dimensional analysis indicated that length/width ratio was unaffected by loss of *lama1* in our small sampling of cells, our 3-dimensional analysis demonstrates that loss of *lama1* or loss of both *lama1* and *pard3* leads to diminished length/width ratio (Figure 5F-F’’; wild type 2.08±0.50; *lama1^−/−^* 2.03±0.55; *lama1^−/−^;MZpard3^−/−^* 1.97±0.39). The difference between wild type and lama1 mutants appears subtle but is significant, likely due to the large numbers of cells measured. Finally, we assayed retinal progenitor cell volume: loss of *lama1* does not affect retinal progenitor cell volume, however, *lama1;MZpard3* double mutant cells are larger than either wild type or *lama1* single mutant (Figure 5G-G’’; wild type 331.97±183.42 μm^3^; *lama1^−/−^* 333.86±201.88 μm^3^; *lama1^−/−^;MZpard3^−/−^* 369.94±229.80 μm^3^).

Taken together, these measurements are a rich source of quantitative information from which to draw a number of conclusions. First, there is variability in the *lama1* mutant misorientation phenotype. We had previously observed this, but did not have a way to quantify it. Mutant embryos can display varying degrees of tissue disorganization, potentially due to the degree to which the cells might self-organize (possibly influenced by aberrant localization of apical polarity complexes) in the absence of extrinsic polarity cues from laminin. Second, although cell size and shape are slightly different in the *lama1* single mutant compared to wild type, change in cell morphology is unlikely to be the cause of the misorientation phenotype. In contrast, the *lama1;MZpard3* double mutant has larger, less elongated cells (greater volume, diminished length/width ratio). Finally, and importantly, at the population level, retinal progenitor cells in *lama1* single mutants and *lama1;MZpard3* double mutants are dramatically misoriented compared to wild type. Despite the appearance of partial rescue in a single optical section (Figure 5A-A’’), these data clearly demonstrate that loss of *pard3* does not rescue the tissue disorganization phenotype in the *lama1* mutant. These 3-dimensional visualization and quantitative analyses underscore the utility of our approach.

## Discussion

A key part of epithelial organogenesis is the establishment of tissue-specific structures which are crucial for eventual organ function. Within these epithelial tissues, cells take on a stereotypical 3-dimensional organization. Much work has gone into identifying molecular signals and pathways that influence this organization. The vertebrate eye is a somewhat unique structure, in which the hemispherical retinal epithelium enwraps the lens. We originally set out to determine how changes in one such class of molecules, the extracellular matrix, affect tissue organization: previously, we found that loss of *lama1* leads to disruptions to tissue polarity and apparent disorganization of the retinal epithelium. 2-dimensional analysis of a limited sampling of cells suggested that cell length was shorter, but length-width ratio seemed unaffected. In addition, loss of *lama1* resulted in ectopic localization of the apical determinant pard3 and other apical markers, and we wondered whether loss of pard3 could rescue these phenotypes. Due to limitations in our ability to visualize and quantitatively analyze 3-dimensional cell shape and orientation, we were not able to test this until now.

LongAxis allows us to take volume data (e.g. confocal z-stacks that are simple to acquire for zebrafish embryos), and run it through a 3-dimensional cell segmentation and analysis pipeline, which includes manual cell selections and filtering based on parameters derived from the manually selected cells. After filtering, LongAxis provides intuitive visualization (in the form of the “urchin plot”) and quantitative analysis of cell orientation and a number of cell morphology descriptors, including length, width, length-width ratio, and volume. The power of LongAxis is in the ability to analyze cell shape and organization at the population level – thousands of cells per eye – rather than manually measuring 2-dimensional features on a limited sampling of cells. This allows us to examine distributions within the cell population as well as variability between individual embryos.

Using LongAxis, we validated our pipeline via manual validation of segmentation and filtering, finding that filtering preferentially removes poorly and incompletely segmented cells. Next, given that vertebrate embryonic development is not deterministic and that variability exists between embryos of the same genotype, we compared results between wild type embryos, finding that LongAxis indeed allows us to detect differences between embryos of the same genotype. Therefore, analysis of multiple embryos of the same genotype is necessary to provide a complete quantitative picture of the phenotype range encompassed.

Finally, we returned to the biological question we initially sought to answer. *lama1* mutant optic cups are comprised of misoriented retinal progenitors which are shorter and slightly less elongated than their wild type counterparts. We had not previously detected the elongation defect, likely due to combination of 2-dimensional analysis and small sample size. Misoriented retinal progenitors appear to cluster together in domains of the optic cup, rather than being scattered throughout the tissue randomly. We speculate that this is due to the ability of cells to self-organize in the absence of extrinsic polarity cues. We did indeed detect and were able to quantify variability between individual *lama1* mutant embryos, with one embryo exhibiting less disruption to cell orientation than the other two. Did removal of the apical determinant *pard3* rescue these phenotypes? Although certain single optical sections looked as though cells were well-oriented toward the lens, 3-dimensional urchin plots demonstrate that cell orientation in *lama1;MZpard3* double mutant optic cups is clearly not rescued; again, misoriented cells cluster together in domains of the optic cup. Further, we detected a change in cell size and shape in the double mutants: cells are larger and less elongated than their wild type or *lama1* single mutant counterparts. The underlying cause of this change in cell size is unknown, but *pard3* has been linked to regulation of proliferation in some systems (Costa et al., 2008); these mechanisms will be interesting to explore moving forward.

LongAxis is currently optimized for the zebrafish optic cup, but could be modified for other epithelial organs. Tissue organization is a crucial aspect of the development of numerous other organs, including brain, ear, and gut. LongAxis is already potentially capable of cell morphology quantification in these other systems; by modifying the code provided, cell orientation analysis could be adapted for a different specific 3-dimensional structure of interest: for example, the convergence point in the eye could be modified to be the midline plane in the developing brain.

As imaging technologies and approaches continue to improve our ability to visualize the cellular basis of tissue assembly and morphogenesis, it is important that our analysis methods also evolve to take advantage of this rich source of 3-dimensional quantitative information.

Tools such as LongAxis will help us connect molecular genetics to cell biology to uncover the mechanisms underlying morphogenesis and development of the visual system and other organs of interest.

## Experimental Procedures

### Zebrafish husbandry and mutant/transgenic lines

All zebrafish husbandry (*Danio rerio*) was performed under standard care conditions in accordance with University of Utah Institutional Animal Care and Use Committee (IACUC) Protocol approval (Protocol #18-02006). Embryos were raised at 28.5-30°C and staged according to time post fertilization and morphology (Kimmel et al., 1995). Mutant lines were previously described: *lama1^UW1^* (Bryan et al., 2016; Semina et al., 2006); *pard3^fh305^* (Blasky et al., 2014). In all cases, maternal-zygotic *pard3* mutants (*MZpard3*) were used.

#### *lama1^UW1^* genotyping protocol

A dCAPS strategy (Neff et al., 1998) was used with the following primers: 5’ GCAGATGCAGCAACCACAGCCAGTCATGTGACCTGCACACCGGCCAACACCT; 3’ GGCTTTCCCCCTCTGATGACACGTAC. PCR annealing temperature, 58°. PCR products were digested with DraIII, which cuts WT (231+47 bp), not mutant (278 bp). Digest products were run on 3.2% Metaphor or 1% Metaphor/1% agarose gel.

#### *pard3^fh305^* genotyping protocol

A CAPS strategy was used with the following primers: 5’ ATTGGCTTCAGCAGTTTTAAGAAA; 3’ ATGATTGGCACTGAGTGAAGAAC. PCR annealing temperature, 61°. PCR products were digested with HpyCH4IV, which cuts mutant (87+68 bp), not WT (155 bp). Digest products were run on 3.2% Metaphor or 1% Metaphor/1% agarose gel.

### RNA synthesis and injections

Capped RNA was synthesized using a pCS2 template (pCS2-EGFP-CAAX) and the mMessage mMachine SP6 kit (Ambion). RNA was purified (Qiagen RNeasy Mini Kit) and ethanol precipitated. 150 pg RNA was injected into the cell of 1-cell embryos.

### Imaging

Embryos were dechorionated at 24 hpf and embedded in 1.6% low melting point agarose (in E2+gentamycin) in Delta T dishes (Bioptechs (#0420041500C)). Images were acquired using a Zeiss LSM710 or LSM880 laser scanning confocal microscope. E2+gentamycin was overlaid, and the dish covered to prevent evaporation. All imaging was performed with a 40X water-immersion objective (1.1 NA). Datasets were acquired with the following parameters: 1024×1024; 0.21 x 0.21 x 0.42 μm voxel size. The entire depth of the optic cup was imaged, resulting in z-stacks of 340-480 slices. All imaging was of live embryos, to avoid distortions that accompany tissue fixation.

### LongAxis MATLAB code

The full LongAxis MATLAB code is available here with annotations: www.kwan-lab.org/longaxis

### LongAxis Segmentation Validation

Segmentation accuracy for all cells was scored manually, by selecting a subvolume, usually containing 50-70 cells. Each cell in the subvolume was examined individually against xy/xz/yz cutaways of the original image data to determine how well the segmentation matched the image data. Accuracy was scored on a scale of 1-5, with 1-4 corresponding to how well the segmentation matched the image data (1 = 90-100%; 2 = 70-90%; 3 = 50-70%; 4 = <50%), and a score of 5 representing unsuccessful segmentation resulting in fused cells.

### Plots

Density, violin (with box and whisker), and stacked bar plots were generated using the ggplot2 package in R. Exponential decay equations were derived and plotted in R using the self-starting asymptotic regression function (SSasymp). Exponential decay equations followed the formula: *y(t)* = *y_f_* + *(y_0_-y_f_)e*^-*λt*, where *y* is the number of excluded cells; *y* starts at *y_0_* and decays towards *y_f_* at rate *λ*.

### Statistics

For comparisons of length, length-width ratio, and volume between wild type, *lama1* single mutant, and *lama1;MZpard3* double mutant, data were compared using ANOVA, followed by Tukey’s test. For comparisons of distributions of angles of deviation, a two-sample Kolmogorov-Smirnov test was carried out in R.

## Acknowledgments

We are grateful to Bruce Appel for providing the *pard3^fh305^* mutant line, and the University of Utah Centralized Zebrafish Animal Resource for zebrafish husbandry. Thanks to members of the Kwan lab for useful discussions and critical reading of the manuscript. This work was supported by grants from the NEI/NIH (R01 EY025378, R01 EY025780) to K.M.K. C.D.B. was supported by the University of Utah Developmental Biology Training Grant (NIH T32 HD007491).

---

**Figure S1.**
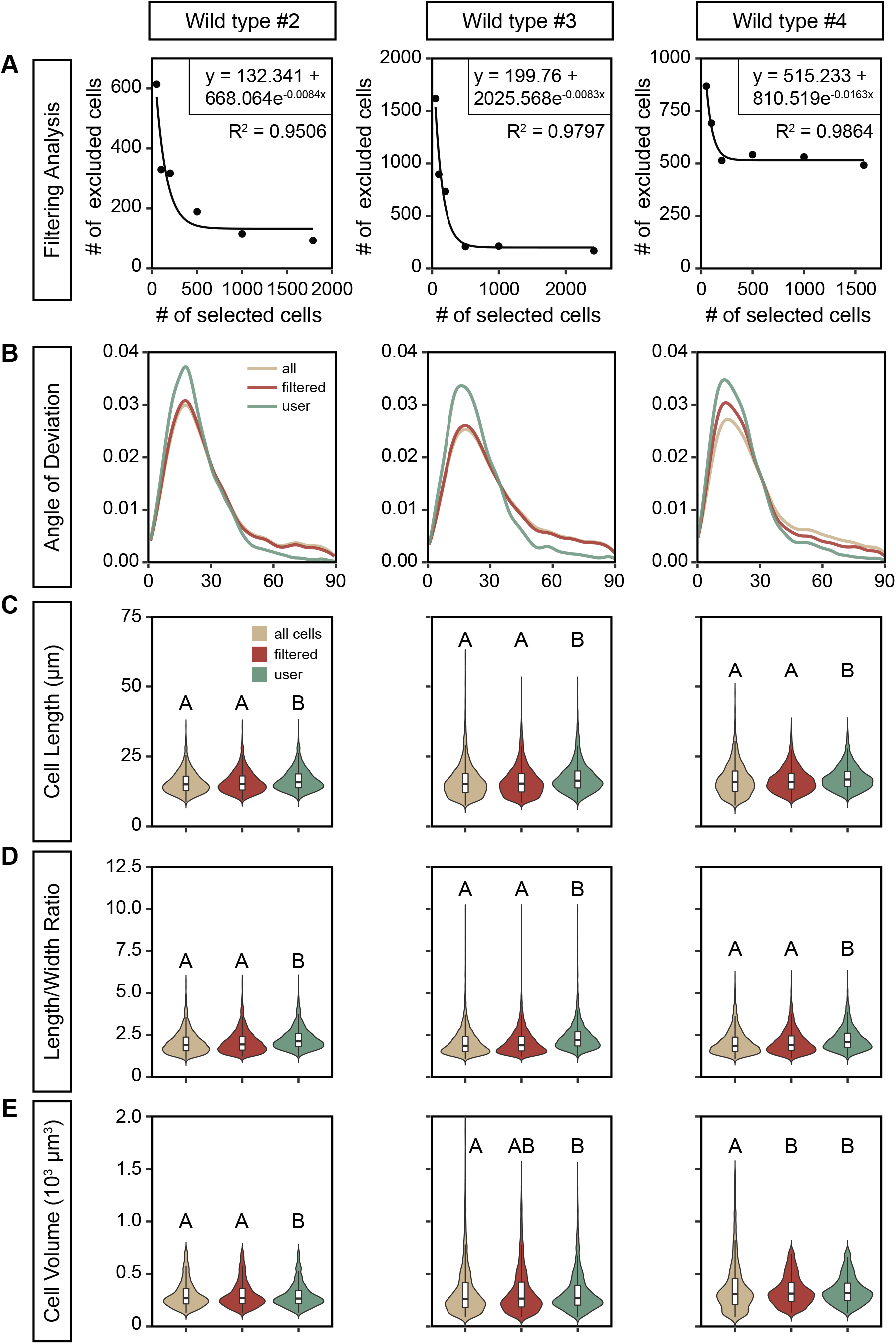

**Figure S2.**
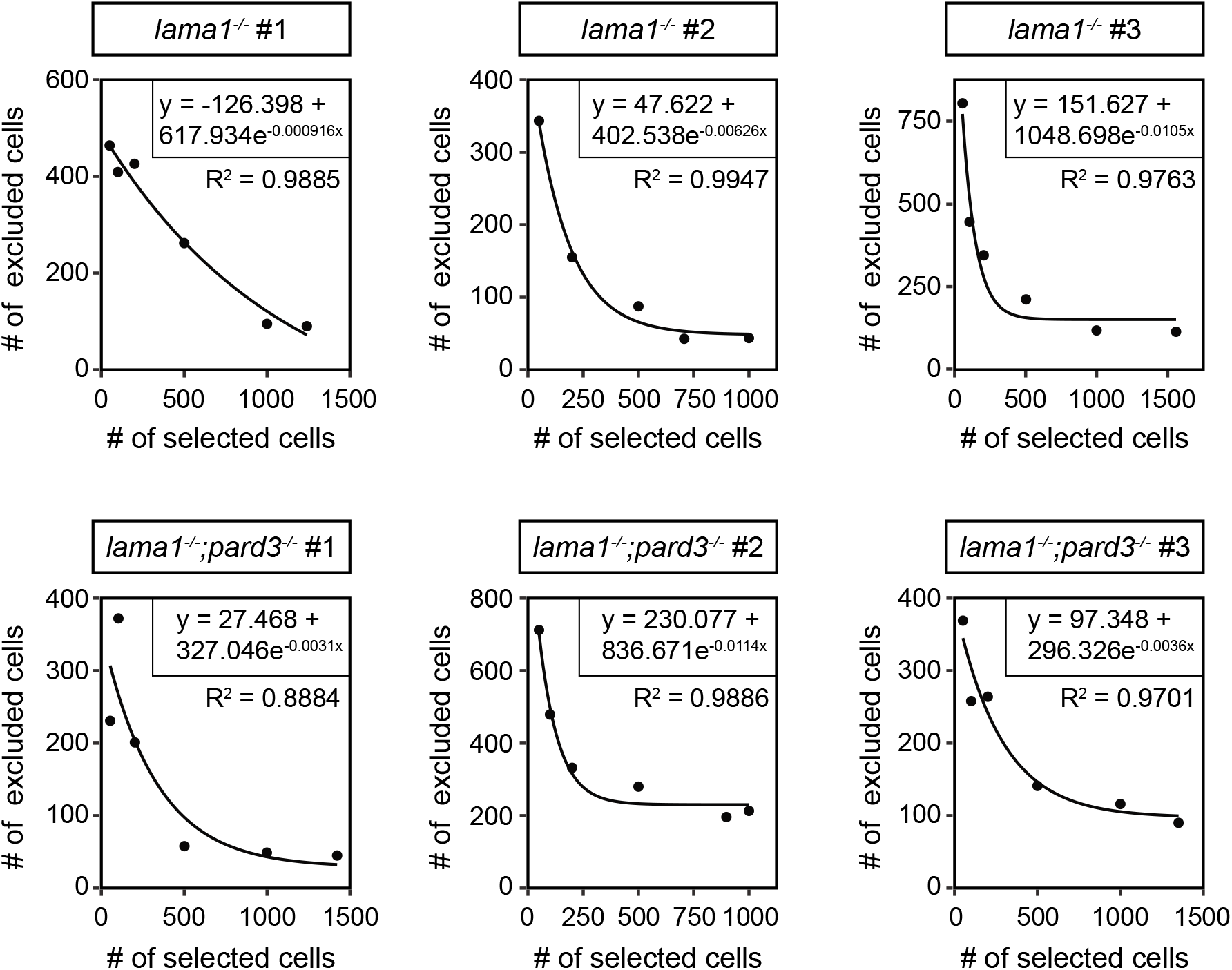

